# Logistic PCA explains differences between genome-scale metabolic models in terms of metabolic pathways

**DOI:** 10.1101/2023.11.27.568810

**Authors:** Leopold Zehetner, Diana Széliová, Barbara Kraus, Juan A. Hernandez Bort, Jürgen Zanghellini

## Abstract

Genome-scale metabolic models (GSMMs) offer a holistic view of biochemical reaction networks, enabling in-depth analyses of metabolism across species and tissues in multiple conditions. However, comparing GSMMs against each other poses challenges as current dimensionality reduction algorithms or clustering methods lack mechanistic interpretability, and often rely on subjective assumptions. Here, we propose a new approach utilizing logisitic principal component analysis (LPCA) that efficiently clusters GSMMs while singling out mechanistic differences in terms of reactions and pathways that drive the categorization.

We applied LPCA to multiple diverse datasets, including GSMMs of 222 *Escherichia*-strains, 343 budding yeasts (*Saccharomycotina*), 80 human tissues, and 2943 *Firmicutes* strains. Our findings demonstrate LPCA’s effectiveness in preserving microbial phylogenetic relationships and discerning human tissue-specific metabolic profiles, exhibiting comparable performance to traditional methods like t-distributed stochastic neighborhood embedding (t-SNE) and Jaccard coefficients. Moreover, the subsystems and associated reactions identified by LPCA align with existing knowledge, underscoring its reliability in dissecting GSMMs and uncovering the underlying drivers of separation.

**Author’s summary:** Genome-scale metabolic models (GSMMs) are comprehensive representations of all the biochemical reactions that occur within an organism, enabling insights into cellular processes. Our study introduces logisitic principal component analysis (LPCA) to explore and compare these biochemical networks across different species and tissues only based on the presence or absence of reactions, summarized in a binary matrix. LPCA analyzes these binary matrices of specific biochemical reactions, identifying significant differences and similarities. We applied LPCA to a range of datasets, including bacterial strains, fungi, and human tissues. Our findings demonstrate LPCA’s effectiveness in distinguishing microbial phylogenetic relationships and discerning tissue-specific profiles in humans. LPCA also offers precise information on the biochemical drivers of these differences, contributing to a deeper understanding of metabolic subsystems. This research showcases LPCA as a valuable method for examining the complex interplay of reactions within GSMMs, offering insights that could support further scientific investigation into metabolic processes.

## Introduction

Genome-scale metabolic models (GSMMs) are mathematical representations of species- or context-specific metabolic reaction networks [1] and have been successfully applied to study diseases [2–4], optimize bioprocesses [5–8], or investigate metabolic differences across species [9–11]. To compare different GSMMs, dimensionality reduction techniques can be applied to the results of simulations, such as flux balance analysis [12]. The flux distributions or growth rates can be predicted for different environmental conditions and serve as input for principal component analysis (PCA). PCA of growth rates was used to suggest potential auxotrophies [9, 11]. While this approach has become a well established method for comparing GSMMs, it necessitates the incorporation of preassumed environmental data. Another challenge is that the predicted growth rates can be similar even in different environments, which might lead to a reduced discriminative capacity. One potential remediation is to focus solely on simulated growth rates from selected environmental conditions exhibiting significant variation across GSMMs; however, this mandates the integration of subjective parameters or thresholds.

Alternative methods analyze binary matrices derived from the presence or absence of reactions in the models. Jaccard coefficients [13] are often utilized to measure similarity between GSMMs, visualized through heatmaps to identify clusters of similar models. However, this method lacks insight into specific pathways driving heterogeneity.

t-distributed stochastic neighborhood embedding (t-SNE) analysis [10, 14], while effective in clustering GSMMs consistently, requires specific prerequisites, such as distance metrics, and hyperparameters, which might result in erroneous clustering outcomes [15–17], and may lack reproducibility due to its non-deterministic nature. Additionally, it does not provide a straightforward identification of key variables driving clustering. In contrast, PCA allows the calculation of loadings representing reactions contributing most to principal components. This enables further analysis by grouping loadings based on gene ontology terms, pathways, or subsystems. However, PCA’s direct application to binary data is not suitable [18] because it relies on variance and covariance calculations, assumptions of continuous distribution, and linear relationships, which are better suited for continuous datasets rather than binary datasets [19]. Thus, as GSMMs become increasingly complex, the need for automated tools to identify the underlying reactions and pathways driving clustering grows, making simultaneous comparison and factor identification essential.

To address the limitations of existing methods, we use logisitic principal component analysis (LPCA) for classifying GSMMs (see Fig 1). LPCA is an adaption of classic PCA to analyze heterogeneity in binary data [18, 20]. In a classic PCA, continuous data is transformed to a new coordinate system such that the greatest variance lies on the first coordinate. Landgraf and Lee introduced LPCA [21], which extends traditional PCA to handle binary data. They reinterpret PCA as a method for projecting data into a lower-dimensional space while maintaining proximity to the original data. For Gaussian data, this involves a straightforward projection, and minimizing squared error. However, for binary data, LPCA projects natural parameters derived from a Bernoulli model while minimizing Bernoulli deviance.

**Fig 1.**
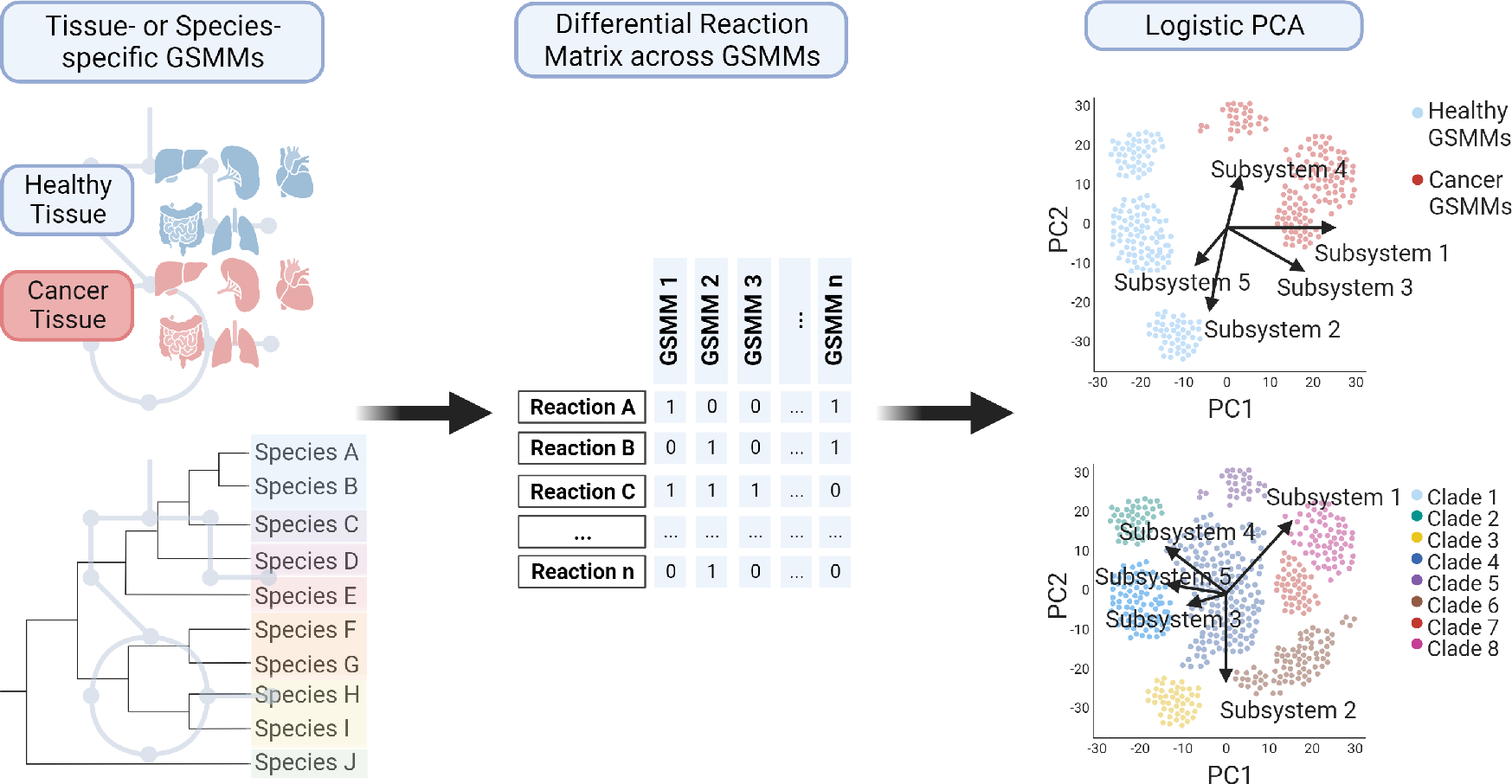
Schematic workflow of applying LPCA to binary reaction matrices derived from GSMMs. (Created with BioRender.com)

LPCA has been used with other biological data, such as binary genomics data [18], but not for the comparison of GSMMs. Here we show that LPCA enables efficient clustering based solely on the presence or absence of reactions. Additionally, using LPCA, reactions were identified as contributing most to clustering, demonstrating its advantages over existing methods like t-SNE and Jaccard coefficients. Furthermore, we identify key subsystems that differentiate GSMMs clusters, a feature not achievable with current methods. We validate our approach by reconstructing phylogenetic associations. Overall, we provide an alternative for streamlined subsystem analysis that elucidates variations across GSMMs.

## Methods

### LPCA on binary reaction matrices

Unless otherwise stated, LPCA was performed on the differential binary pan-reaction matrix, Δ***X***, using the logisticPCA function from the “logisticPCA” package (v0.2) [21] in R (v4.3.1). Due to the large dimensions of the binary matrices, partial decomposition was chosen, as recommended in the “logisticPCA” package documentation. This approach involves computing only a few eigenvalues instead of all of them, thereby speeding up the computation process.

The binary data matrix Δ***X*** (*N × R*) consists of *N* rows, where every row represents one context- or species-specific GSMM, and *R* columns represent the reactions. LPCA minimizes the Bernoulli deviance *D*, which quantifies the difference between the binary data matrix Δ***X***, and the LPCA reconstruction **Θ**, representing the estimated natural parameters of the Bernoulli model,

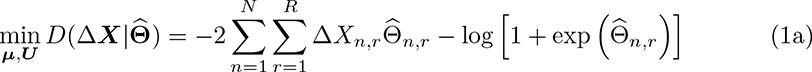

with

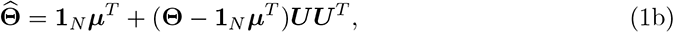

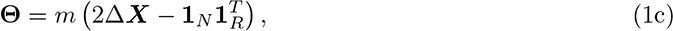

with a tunable parameter *m* that, in default mode, is automatically selected by the logisticPCA function. **Θ** (*N × R*) denotes the matrix of natural parameters from the saturated model. A saturated model in the context of binary data and Bernoulli distributions is one in which the estimated probabilities exactly match the observed data Δ***X***. The results from the logisticPCA function include the mean parameter vector ***µ*** (*R ×* **1**) and the loading matrix ***U*** (*R × i*), where *i* belongs to the number of principal components. ***U*** and ***µ*** are solved such that the Bernoulli deviance *D* is minimized.

The principal component scores ***S*** (*N × i*) are then obtained from ***U*** and ***µ*** within the logisticPCA by:

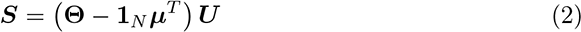

The set of loadings ***U_∗,r_*** across all principal components *i* for a reaction *r* represents a loading vector.

We refer to a loading vector containing loadings for the first two principal components as reaction-centric loading vector. These reaction-centric loading vectors can be added to LPCA score plots as arrows to visualize how each reaction contributes to the clustering.

To avoid overwhelming complexity in visualization, we introduce subsystem-centric loading vectors as proxies. For each principal component *i*, we compute the average loading across all reactions *r* within subsystem *j*, denoted as

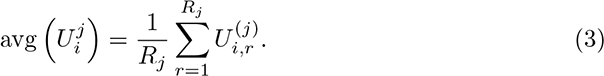

Here, *R_j_*is the number of reactions in subsystem *j*, and *U* ^(*j*)^ is the loading of principal component *i* for reaction *r* within the subsystem *j*.

In Fig 2, 4, 5, and 7 we visualize these average loading vectors for the first two principal components.

**Fig 2.**
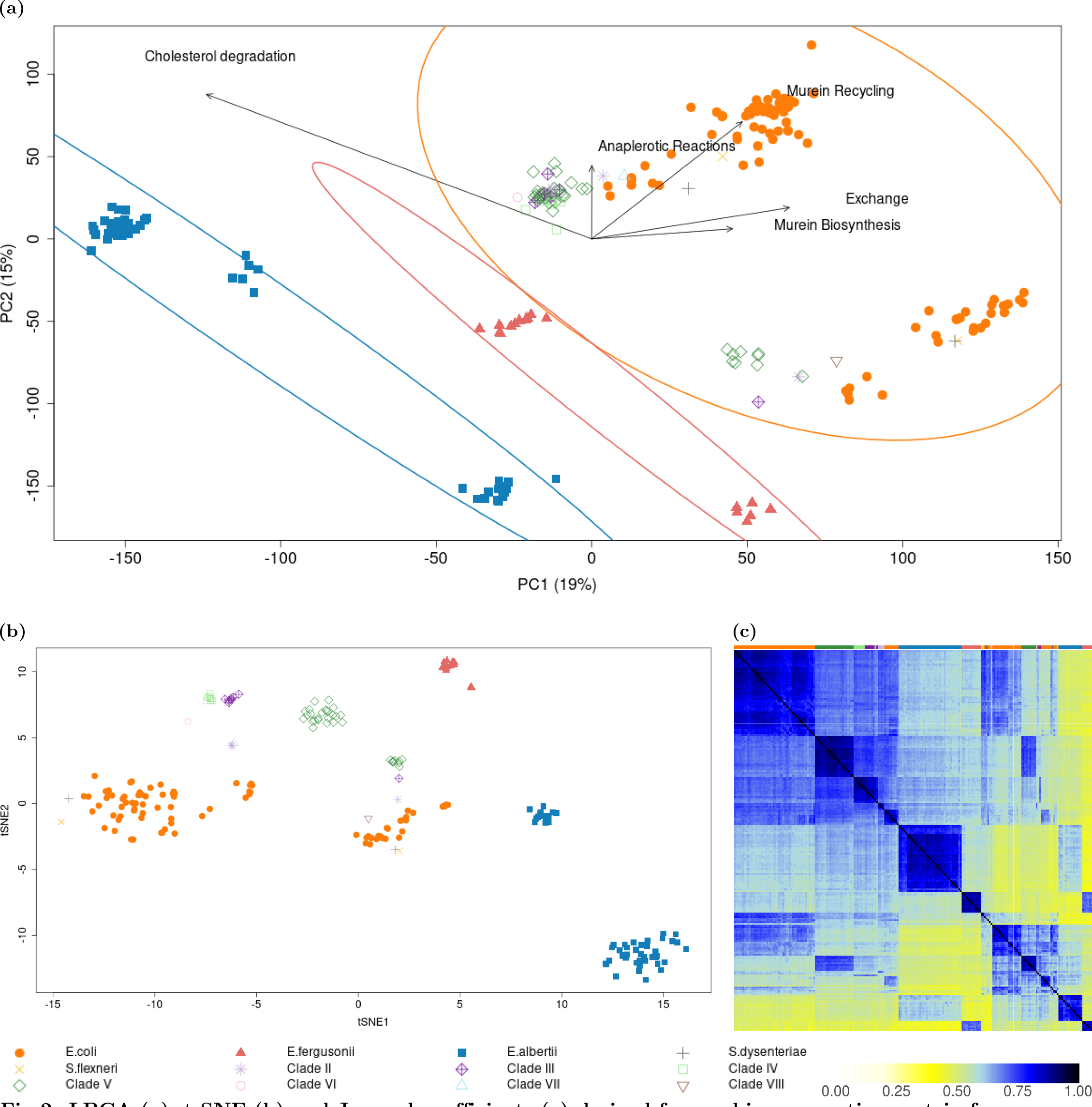
LPCA (a), t-SNE (b) and Jaccard coefficients (c) derived from a binary reaction matrix from differential reactions in. 222 *Escherichia* GSMMs. In panels (a) and (b), points represent individual GSMMs, with different genera indicated by distinct symbols and colors. The top row in panel (c) uses these same colors to indicate the corresponding genera. Circles in panel (a) highlight clusters of *E. albertii* strains (blue), *E. fergusonii* strains (red), and a mixed cluster of *E. coli*, *S. dysenteriae*, *S. flexneri*, and Clades II to VIII (orange). Labeled arrows in panel (a) denote subsystem-centric loading vectors from LPCA (refer to the results and methods section for definitions)

To rank the contributions of the subsystems to the principal components, we compute the Euclidian norm of the subsystem-centric loading vectors across the first two principal components

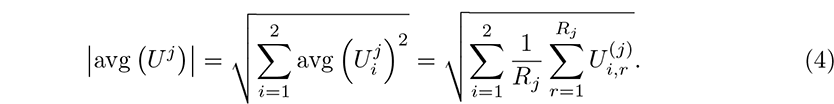

As an alternative ranking measure, we first calculate the Euclidian norm of every reaction-centric loading vector, and then the average as

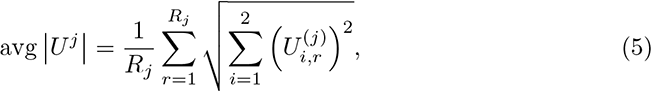

where the inner sum traverses over the first two principal components.

Both measures were employed to identify key subsystems responsible for driving the observed differentiation.

### Data sources, collection and pre-processing

#### Differential binary pan-reaction matrix, **Δ***X*

We acquired GSMMs and metabolic reconstructions from four distinct sources as outlined below. For each dataset, we generated a binary “pan-reaction” matrix ***X*** *∈ {*0, 1*}^N×R^*, where each of the *N* rows corresponds to a GSMM and each of the *R* columns represents a reaction, using COBRApy (v0.26.3) [22]. These matrices indicate whether individual reactions are present (1) or absent (0) within species-specific GSMMs or context-specific reconstructions and provide a comprehensive overview of the reactions present across all the GSMMs. To simplify the matrices, we removed columns containing only 1 or 0. We refer to the resulting data as the differential binary reaction matrix Δ***X*** of the pan-GSMM or the pan-reconstruction.

- ***Escherichia*** . We used 222 GSMMs of *Escherichia* species reconstructed by Monk [9], grown across 570 environmental conditions. The resulting pan-GSMM contains 3342 reactions, each present in at least one strain. Of these, 1688 reactions are not consistently present across all strains, which thus form the differential matrix Δ***X****_Escherichia_ ∈ {*0, 1*}*^222^*^×^*^1688^.

During our analysis, we discovered that the reaction labeled as “PRCOA1” is associated with “Cholesterol degradation” [23] rather than its initial association with “Histidine Metabolism” [9]. Consequently, we established a new subsystem named “Cholesterol degradation”, reassigning “PRCOA1” and related reactions based on BiGG annotations. Subsequently, we recalculated the LPCA and subsystem-centric loadings to reflect this adjustment.

- ***Firmicutes***. We used 2943 GSMMs of the phylum *Firmicutes* from the Agora2 dataset [14] forming a differential matrix Δ***X****_Firmicutes_ ∈ {*0, 1*}*^2943^*^×^*^5267^.
- ***Fungi*** . We used 343 GSMMs of fungi reconstructed by Lu et al. [10] based on previous sequencing data [24]. The resulting pan-GSMM contains 4599 reactions, each present in at least one strain. Of these, 2519 reactions are not consistently present across all strains, forming the differential matrix Δ***X****_Fungi_ ∈ {*0, 1*}*^343^*^×^*^2519^.
- ***Human*** . We used normalized gene expression data from 50 healthy tissue samples [25] and 30 cancer tissue samples [26] sourced from the Human Protein Atlas [25, 26] to generate context-specific genome-scale metabolic reconstructions. Reactions were taken from the latest iteration of the human metabolic model, “Human1” [27], with gene expression considered for normalized transcripts per million (nTPM) values exceeding 0.2. To avoid bias from gap-filling, only reactions with gene assignments were included in the context-specific reconstruction. The resulting pan-reconstructions contained 7975 reactions. Of these, 2390 reactions are not consistently present across all reconstructions, which thus form the differential matrix Δ***X***_Human_ *∈ {*1, 0*}*^80^*^×^*^2390^.

### Additional computational Analyses

#### t-SNE on binary reaction matrices

Binary reaction datasets from context-specific GSMMs were analyzed by Hamming-distance based t-SNE calculation using the R package “Rtsne” (v0.16) [28]. For plotting, only the first two t-SNE values were considered.

#### Jaccard coefficients on binary reaction matrices [29]

Jaccard coefficients were calculated based on binary reaction pairwise between context-specific GSMMs using:

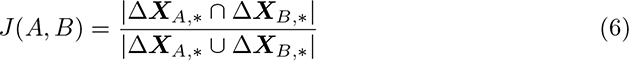

where *J* (*A, B*) is the Jaccard coefficient between differential reactions of GSMMs *A* and *B*, consisting of reactions Δ***X****_A,∗_*, and Δ***X****_B,∗_*, respectively. *|*Δ***X****_A,∗_ ∩* Δ***X****_B,∗_|* is the intersection of reactions between GSMMs *A* and *B*, and *|*Δ***X****_A,∗_ ∪* Δ***X****_B,∗_|* is union of reactions between GSMMs *A* and *B*.

#### Principal component analysis

##### Escherichia

For visual comparison, the PCA plot based on simulated growth rates from environmental conditions was reproduced for *Escherichia* strains from [9]. To obtain a similar clustering, a correlation matrix was computed, and PCA was performed using the princomp function in R.

### Phylogenetic analysis

#### Escherichia

Proteins from genomic sequences (downloaded from Enterobase - v1.1.5 [30]) were predicted using “prodigal” (v2.6.3) [31], followed by phylogenetic comparison using “OrthoFinder” (v2.5.5) [32]. The phylogenetic tree was then plotted using the “ape” package (v5.7-1) [33] in R.

#### Fungi

The phylogenetic tree was reproduced using the “ape” package [33] based on a Newick file published by Shen *et al* [24].

### Cophenetic correlation coefficient

We used the cophenetic correlation coefficient [34] to examine similarity between LPCA scores, and phylogenetic trees, where possible. Initially, we computed the pairwise distances among LPCA scores using both Euclidean and Manhattan metrics. These distances were then subjected to hierarchical clustering, resulting in the formation of dendrograms. The obtained dendrograms were compared using the cophenetic correlation coefficient by applying the cophenetic, and cor functions in R. Due to the inherent non-deterministic nature of t-SNE, a comparison based on the cophenetic correlation coefficient is not meaningful.

### Subsystem analysis using multinomial logistic regression

A multinomial logistic regression (MLR) model, using the multinom function (“nnet”, v7.3-19), was applied to calculate the contribution of specific reactions to pre-defined clusters of GSMMs (i.e. phylogenetic clades/tissue types). Parameter estimates were derived for each reaction, reflecting their relative contributions to the cluster membership. To quantify the aggregate influence of reactions within biological subsystems, we calculated the Euclidean norm of the parameter estimates for each reaction, which provides a measure of the reaction’s importance. These values were then averaged by subsystem, offering a subsystem-level perspective on the factors driving the clustering of metabolic models following:

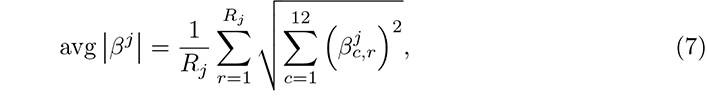

where *β* is the coefficient for reaction *r* in subsystem *j*, for one of the 12 clades *c*. *R_j_* is the number of reactions in subsystem *j*. Before plotting, subsystems were normalized by dividing every subsystem by the highest value from MLR and LPCA.

## Results and Discussion

### Comparison of 222 strain-specific GSMMs from the genus

#### Escherichia

We used LPCA to analyze 222 strain-specific GSMMs of the genus *Escherichia* [9]. Both the differential binary pan-reaction matrix Δ***X*** (Fig 2a) and the full binary pan-reaction matrix (Fig S1) were subjected to LPCA, and showed similar clustering behavior. Overall, seven subclusters were identified, three of which were exclusively associated with *E. albertii* strains (blue in Fig 2a), two with *E. fergusonii* strains (red in Fig 2a, red), and the remaining two consist of *E. coli*, *S. dysenteriae*, *S. flexneri*, as well as Clades II to VIII (orange in Fig 2a). While *E. coli* strains did not segregate from *Shigella* species, they tended to accumulate on one side of the two subclusters within the orange ellipse. In contrast, Clades II to VIII gathered on the opposite side.

### LPCA aligns with established GSMM classification methods and phylogenetic analysis

To validate our LPCA analysis, we compared it with t-SNE analysis (see Fig 2b) and a Jaccard coefficient analysis (see Fig 2c). Overall the three methods showed similar clustering behavior.

- t-SNE analysis found two distinct clusters for *E. albertii* (blue in Fig 2b), contrasting the three clusters identified by LPCA. Additionally, t-SNE identified only one cluster for *E. fergusonii* (red in Fig 2b), as opposed to two clusters observed with LPCA. Both LPCA and t-SNE identified two identical clusters for *E. coli* (orange in Fig 2b). However, t-SNE provided better resolution for the remaining clades compared to LPCA.
- Ten permutations of the sample order resulted in substantial variability in the t-SNE-derived coordinates, highlighting the sensitivity of this method. In contrast, the scores obtained from the LPCA approach remained remarkably stable across the same number of permutations (compare Fig S5).
- Hierarchical clustering of pairwise Jaccard coefficients revealed two distinct clusters for *E. albertii*, two clusters for *E. fergusonii*, and two mixed clusters comprising *E. coli* and the remaining clades (Fig 2c).

Again, clustering patterns remained essentially consistent across all methods when using the full binary pan-reaction matrix instead of the differential matrix (compare Fig 2 with Fig S1).

Next, we tested if the clustering by LPCA aligns with a phylogenetic analysis (Fig S3). We compared LPCA scores with a whole-genome based phylogenetic tree using the cophenetic correlation coefficient [34]. LPCA scores showed a good cophenetic correlation with the phylogenetic tree when Manhattan distances (0.61) were used to calculate the pairwise distance between scores. The correlation weakened when using Euclidean distance. This drop in correlation with Euclidian distance (0.37) is expected as it is more suited for continuous data rather than binary data. Consequently, we suggest that Manhattan distances derived from LPCA scores effectively conserve phylogenetic relations.

Finally, when comparing LPCA to standard PCA using simulated growth rates under different environmental conditions, we observe a better separation of GSMMs, especially between *E. coli* and *E. albertii* (Fig S2).

### LPCA pinpoints key metabolic subsystems distinguishing GSMM clusters

To identify the factors driving the observed clusters, we analyzed subsystem-centric loading vectors derived from LPCA, as described in Equation (2) in the Methods section. These vectors aim to measure the contribution of the entire metabolic subsystem (like glycolysis, etc.) rather than individual reactions. We find that cluster separation was predominantly influenced by (the top) five subsystems: “Murein Biosynthesis”, “Murein Recycling”, “Anaplerotic Reactions”, “Exchange”, and “Histidine Metabolism” (Fig 2a). Notably, when employing a different ranking method based on (4) rather than (3), four out of these five driver subsystems remained consistent (see Tab S1), underscoring their robust impact on the clustering.

The subsystem “Exchange” consists of only one reaction:

Methylmalonate-semialdehyde dehydrogenase (“MMSAD3”). Its presence in this list is unexpected given that the *Escherichia* GSMMs were reconstructed to compare 570 environmental conditions. Thus, the corresponding exchange systems should be universally present in all models, regardless of genetic evidence. According to BiGG annotation, “MMSAD3” is indeed categorized as exchange but is associated with propanoate metabolism according to KEGG. We suspect that “MMSAD3” might have been incorrectly assigned.

“Murein Recycling” and “Murein Biosynthesis” are known to be highly conserved subsystems among *Escherichia* strains, with 88 % of the reactions being shared across all *Escherichia* strains [9]. The highest loading values were obtained for D-alanyl-D-alanine dipeptidase (“ALAALAD” from the “Murein Recycling” subsystem), which is conserved in 82 % *E. coli* s, in 100 % *Shigella*s, and in 2 % Clades II to VIII. In contrast, “ALAALAD” is completely absent in the more distinct clades, *E. albertii* and *E. fergusonii*. D-alanyl-D-alanine dipeptidase (*vanX*) is responsible for cleaving D-alanyl-D-alanine dipeptide. This enzyme allows bacteria to use D-alanine as a carbon source and modify peptidoglycan structures [35]. Intriguingly, modifications in these peptidoglycan precursors, such as the substitution of D-alanyl-D-alanine with D-alanyl-D-lactate, confer resistance to the antibiotic vancomycin, which targets terminal D-alanyl-D-alanine residues to prevent crosslinking [35, 36].

In further exploring the capabilities of LPCA, we analyzed the set of differential reactions within the “Histidine Metabolism” subsystem when contrasting the clades *E. albertii* and *E. coli*, as an example. We noted virtually identical loading values for the reactions “HISDr”, “URCN”, and “IZPN” (Tab S2). These reactions detail the stepwise breakdown of histidine into N-formimidoyl-L-glutamate via urocanate and 4-imidazolone-5-propanoate. This pathway was found to be twice as prevalent in *E. albertii* strains (25 %) compared to *E. coli* strains (12 %). This differential trait suggests that at least certain *E. albertii* strains might possess the metabolic capability to utilize histidine as a source of carbon and nitrogen.

One particular reaction, “PRCOA1”, prevalent in 61 % of *E. albertii* -specific GSMMs, appeared as a significant difference, see Tab S2. However, we found that this reaction might be misallocated to “Histidine Metabolism”. According to the BiGG database [37], “PRCOA1” converts CoA-20-hydroxy-cholest-4-en-3-one C5 side chain to Androst-4-ene-3,17-dione, and belongs to “Cholesterol degradation”, rather than “Histidine Metabolism” (see reaction-centric loadings in Tab S2). Thus, we newly introduced the so far missing subsystem “Cholesterol degradation”, reassigned “PRCOA1” (and associated reactions according to KEGG), and repeated LPCA (see Fig 2a) as well as computing the subsystem loadings new. This time “Cholesterol degradation” replaced “Histidine metabolism” among the top five driving subsystems. The results from Fig 2 suggest that LPCA is capable of distinguishing between phylogenetically distant species by analyzing their sets of differential reactions.

Additionally, the loading values from LPCA help identify the key drivers for the observed separation between species and may also hint at misannotations. Unlike other methods, this identification happens simultaneously with the computation of LPCA scores, offering insights into both the extent of metabolic differences and the specific distinctions driving them.

### MLR reveals subsystems contributing to phylogenetic classification

To validate our interpretation of the subsystem-centric loadings from LPCA, we used a MLR model to independently evaluate the contribution of each reaction to phylogenetic clades (see methods for details). By grouping the reaction-centric parameters by subsystem (see Methods 6), we observed different subsystems to be the most influential drivers for separation in comparison to LPCA-derived subsystem-centric loadings. As displayed in Fig 3, “Pentose Phosphate Pathway”, “Lipopolysaccharide Biosynthesis / Recycling”, “Pentose and Glucuronate Interconversions”, “Murein Biosynthesis”, and “Anaplerotic Reactions” are the five most important subsystems, responsible for the separation of clades. Two subsystems are shared between MLR and LPCA within the top five subsystems: “Murein Biosynthesis”, and “Anaplerotic Reactions”. Moreover, “ALAALAD” is again the most influential reaction within the subsystem “Murein Biosynthesis” in MLR. Additionally, it seems that a lower number of subsystems may impact the clustering, since 8 subsystems have a normalized value above 0.5 in MLR, while only 4 subsystems have a normalized value above 0.5 in LPCA. Normalized values in MLR and LPCA were obtained by dividing every subsystem-centric value by the maximum value. While MLR can pinpoint key determinants of a pre-defined categorization, LPCA provides a notable benefit through the detection of possibly new (sub)clusters. Such subclusters could reveal metabolic distinctions that might be missed when exclusively depending on the primary phylogenetic categorization. Moreover, LPCA allows for the identification of the orientation of reaction- or subsystem-centric loadings, an analysis unachievable with MLR.

**Fig 3.**
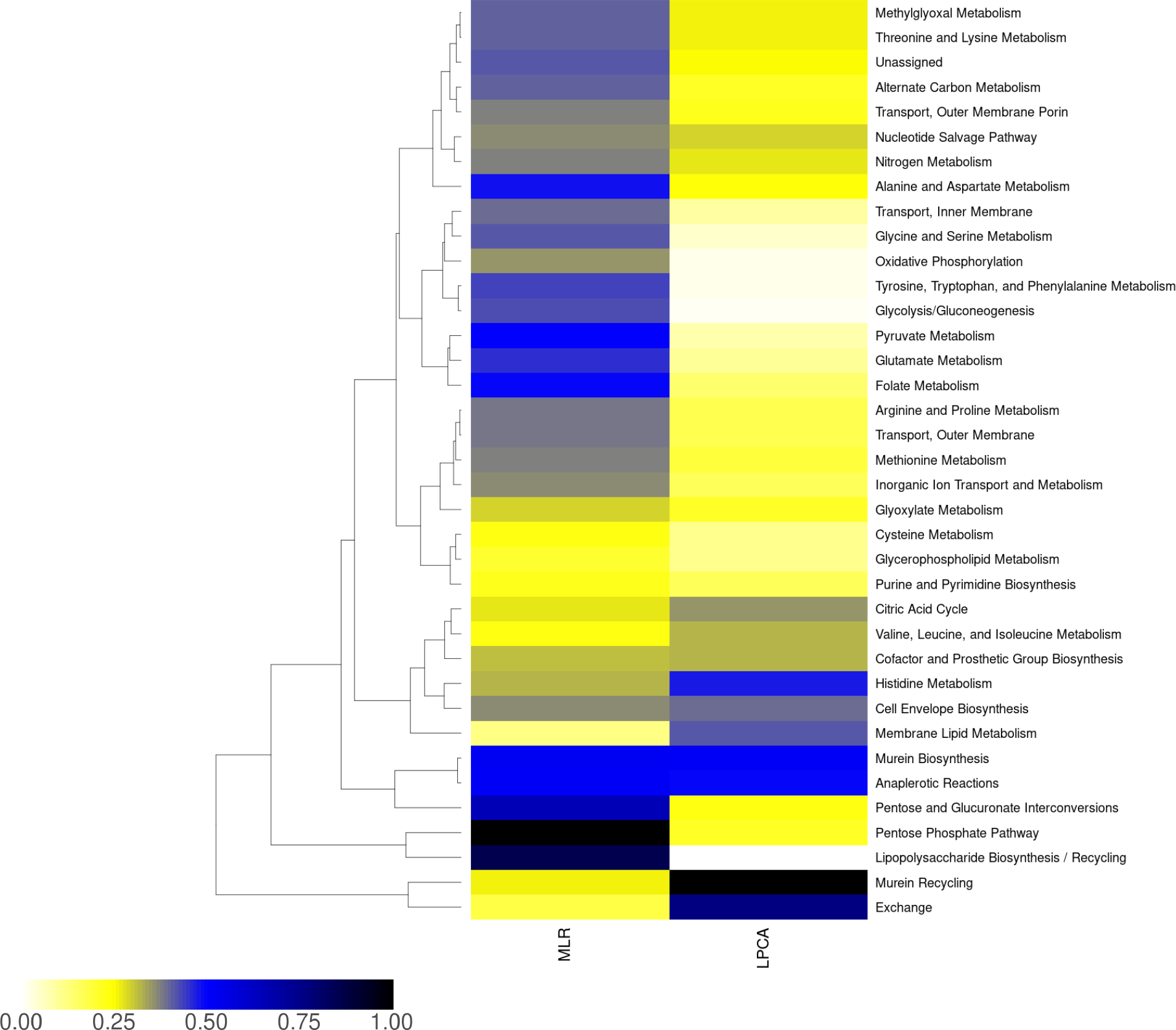
Impact of subsystems derived from LPCA and MLR for *Escherichia* GSMMs. MLR: contribution of subsystems to phylogenetic classification normalized to the maximum value. LPCA: subsystem-centric loadings normalized to maximum loading (refer to methods section for details)

### Comparison of 343 yeast-specific GSMMs from the subphylum Saccharomycotina

We applied LPCA to the set of differential reactions extracted from 343 yeast-specific GSMMs [10], mostly obtained through a recent sequencing effort [24], see 4a. Similar to the LPCA analysis of GSMMs for *Escherichia* (see above), our findings reveal a clustering of species with closer phylogenetic relationships. However, when considering the first two principal components, the separation is less pronounced for yeast, accounting for only 27 % of the variance, in contrast to 34 % for *Escherichia* species. In the phylogenetic tree Fig S6, similar clades were observed to group into closely related clusters as in the LPCA plot Fig 4a. Notably, the *Lipomycetaceae* clade (purple) was distinctly isolated from other clades. In contrast, clades with closer phylogenetic relationships formed subclusters. In particular, the *Saccharomycodaceae*-specific GSMMs (green) constituted a distinct cluster, despite their close phylogenetic relation to *Saccharomycetaceae*.

**Fig 4.**
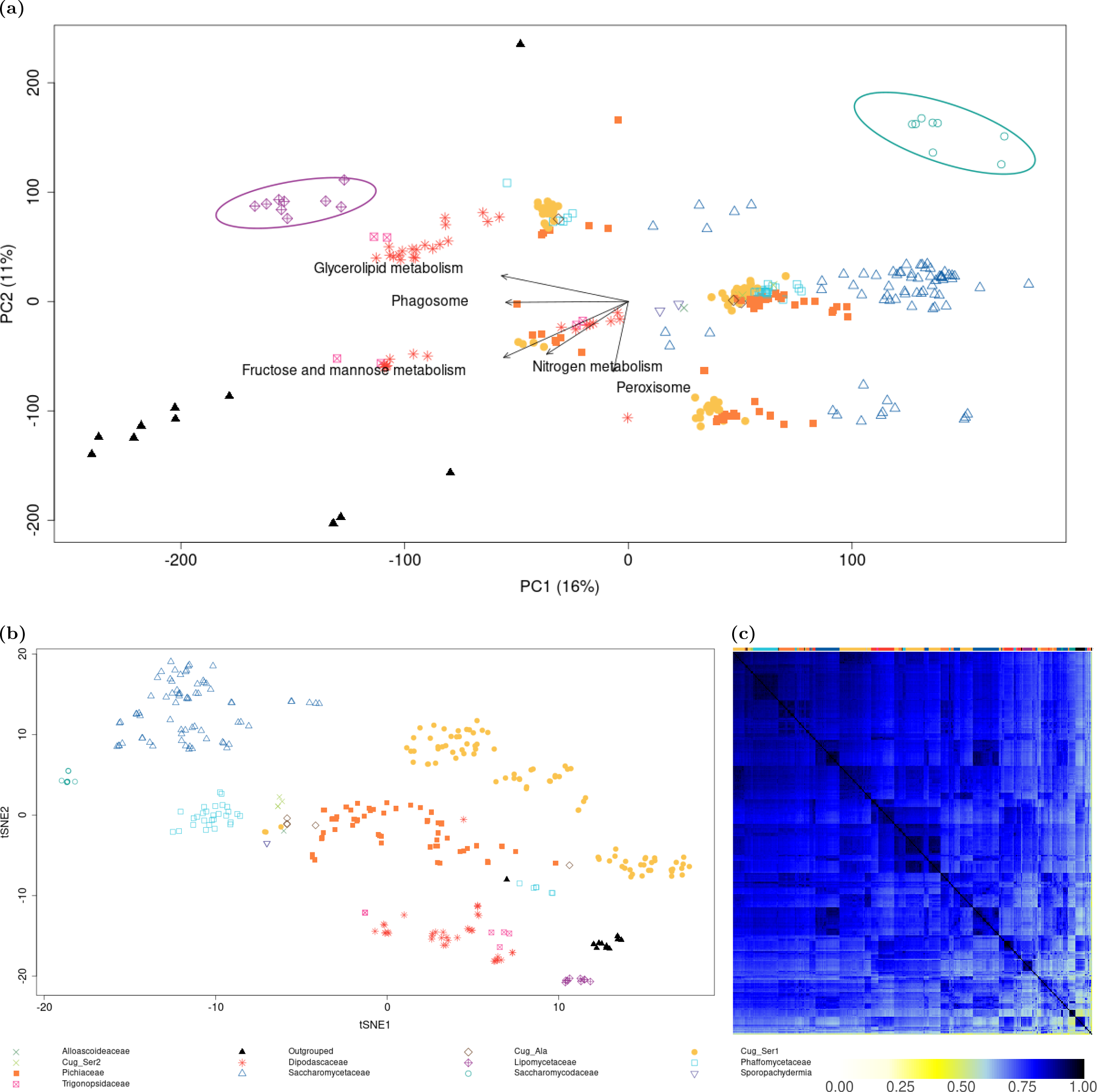
LPCA (a), t-SNE (b), and Jaccard coefficients (c) derived from a binary reaction matrix from yeast-specific GSMMs. In panels (a) and (b), points represent individual GSMMs, with different genera indicated by distinct symbols and colors. The top row in panel (c) uses these same colors to indicate the corresponding genera. Circles in panel (a) highlight a cluster of the *Lipomycetaceae* clade (purple), and the *Saccharomycodaceae* (green). Labeled arrows in panel (a) denote subsystem-centric loading vectors from LPCA (refer to the results and methods section for definitions)

Again, our analysis identified the top five subsystems critically contributing to cluster separation: “Glycerolipid metabolism”, “Phagosome”, “Nitrogen metabolism”, “Fructose and mannose metabolism”, and “Peroxisome”. However, comprehensively examining these pathways in yeast (as we did above for *Escherichia*) faces two major challenges (i) limited knowledge on fungal metabolism – yeast and other fungi remain relatively understudied compared to mammalian and prokaryotic cells [38]; and (ii) a significant proportion of reactions (*>*50 %) are in the subsystem “Unassigned” in the current dataset.

### Comparison of healthy and cancerous tissues

Previous studies have demonstrated the effectiveness of PCA in distinguishing between healthy and cancerous tissues using transcriptomic data [39, 40]. Here, we replicated this finding using normalized gene expression data taken from the Human Protein Atlas [25, 26], see Fig S7. When restricting the analysis to genes annotated within the generic “Human1” reconstruction, the distinction became less evident, see Fig S7.

Further narrowing the analysis to genes derived from the annotations in context-specific GSMMs (see Methods for details) resulted in the loss of differentiation (Fig S7). This outcome may be attributed to the notably fewer genes (561) in these models compared to the broader set in “Human1” (containing 2897 metabolic genes). We explored if LPCA could recover the differentiation between healthy and cancerous tissues using the differential set of reactions in these GSMMs. Indeed, Fig 5a indicates the feasibility of such recovery via LPCA.

**Fig 5.**
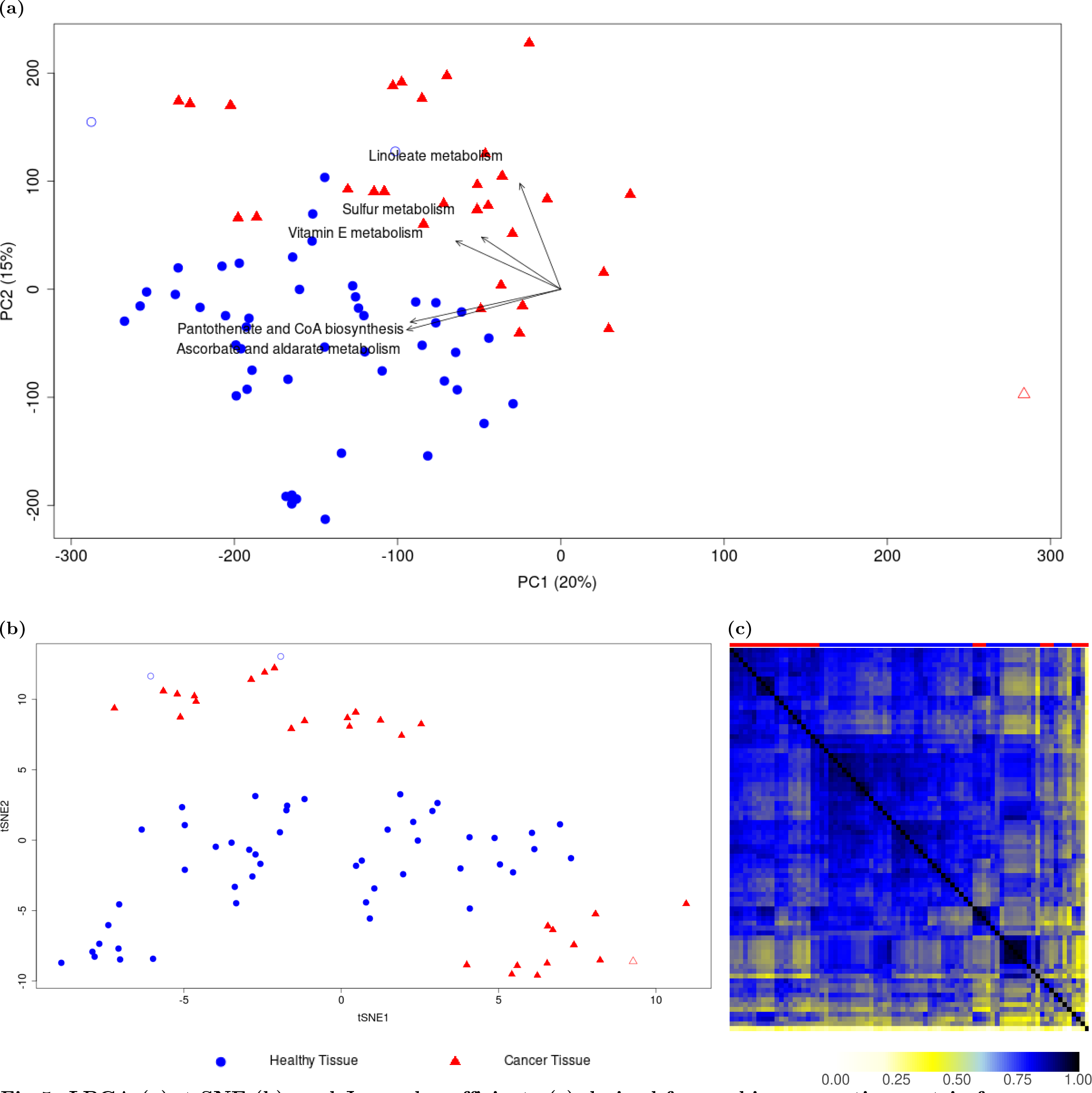
LPCA (a), t-SNE (b), and Jaccard coefficients (c) derived from a binary reaction matrix from context-specific reconstructions from healthy and cancerous tissues. In panels (a) and (b), points represent individual reconstructions, with reconstructions from healthy (blue squares) and cancerous tissues (red triangles). The top row in panel (c) uses these same colors to indicate the corresponding tissue. t-SNE appears less effective in identifying outliers (red open triangle, and blue open circles) compared to LPCA and Jaccard coefficients. Labeled arrows in panel (a) denote subsystem-centric loading vectors from LPCA (refer to the results and methods section for definitions)

Again, in comparison to t-SNE (Fig 5b) and Jaccard coefficients (Fig 5c) suggests that both t-SNE and LPCA can distinguish healthy from cancerous reconstructions. However, t-SNE appears less effective in identifying outliers (Fig 5a and Fig 5b, red, unfilled triangle) compared to LPCA and Jaccard coefficients. In addition, two healthy tissue samples (Fig 5, blue, unfilled circles) tended to cluster with reconstructions from cancer tissues, showing a more pronounced clustering when using LPCA compared to t-SNE. LPCA facilitates a clearer interpretation of the underlying factors, aiding in the identification of differences between context-specific GSMMs.

When analyzing the LPCA-derived subsystem-centric loadings, “Linoleate Metabolism” emerged as a crucial subsystem for distinguishing between context-specific GSMMs. Within this subsystem, the reaction “MAR02438” exhibited the highest loading value and is associated with the genes *PTGS1* or *PTGS2*. These genes play a pivotal role in prostaglandin synthesis and are linked to the vascular endothelial-derived growth factor (VEGF) signaling pathway, critical for angiogenesis [41]. *PTGS2* holds significant importance in cancer research. Inhibitors targeting the enzyme COX2, encoded by the *PTGS2* gene, such as celecoxib, have demonstrated cancer-retarding properties [42].

Another important insight came from the subsystem “Ascorbate and alderate metabolism”, which was found to be top-ranked in subsystem-centric loadings. It consists of only one reaction-centric loading (“MAR08346”), pointing towards healthy tissue samples (Fig 5a), which indicates a higher presence in healthy tissues than cancerous tissues. The reaction describes the reversible conversion of “L-gulonate” to “L-gulono-1,4-lactone” in the endoplasmic reticulum and is catalyzed by the enzyme “Regulacin” (*RGN*). Besides its metabolic role, *RGN* is involved in calcium homeostasis, antioxidant defense, apoptosis, and cell proliferation [43]. Recently, *RGN* has been identified to be downregulated in several cancer cells [44] and that survival of cancer patients is positively correlated with a higher expression level of *RGN* [45].

In the “Human1” GSMM [27], “L-gulonate” can be converted to “L-gulono-1,4-lactone” (*RGN*, endoplasmic reticulum), “glucuronate” (*AKR1A1*, endoplasmic reticulum), or to “3-dehydr-L-gulonate” (*CRYL1*, cytoplasm). High *CRYL1* -expression has been shown to increase survival rate at least in clear cell renal cell carcinoma patients, while *CRYL1* silencing led to increased cell migration and proliferation [46]. In contrast, *AKR1A1* was found to be upregulated in many cancer cells and is associated with drug resistance [47]. Since *AKR1A1* catalyzes the final conversion step from “D-glucose” to “L-gulonate” via “glucuronate”, while *RGN*, and

*CRYL1* are downregulated, an accumulation of “L-gulonate” might emerge in cancerous cells and could be further investigated. This finding is supported by the subsystem analysis using MLR (Fig 6), where the subsystem “Ascorbate and alderate metabolism” is ranked on top as well. Further top-ranked subsystems from MLR include “Terpenoid backbone biosynthesis”, “Metabolism of other amino acids”, “Tricarboxylic acid cycle and glyoxylate/dicarboxylate metabolism”, and “Phosphatidylinositol phosphate metabolism”, none of them shared with the top-ranked subsystems from LPCA, being “Linoleate metabolism”, “Pantothenate and CoA biosynthesis”, “Vitamin E metabolism”, and “Sulfur metabolism”.

**Fig 6.**
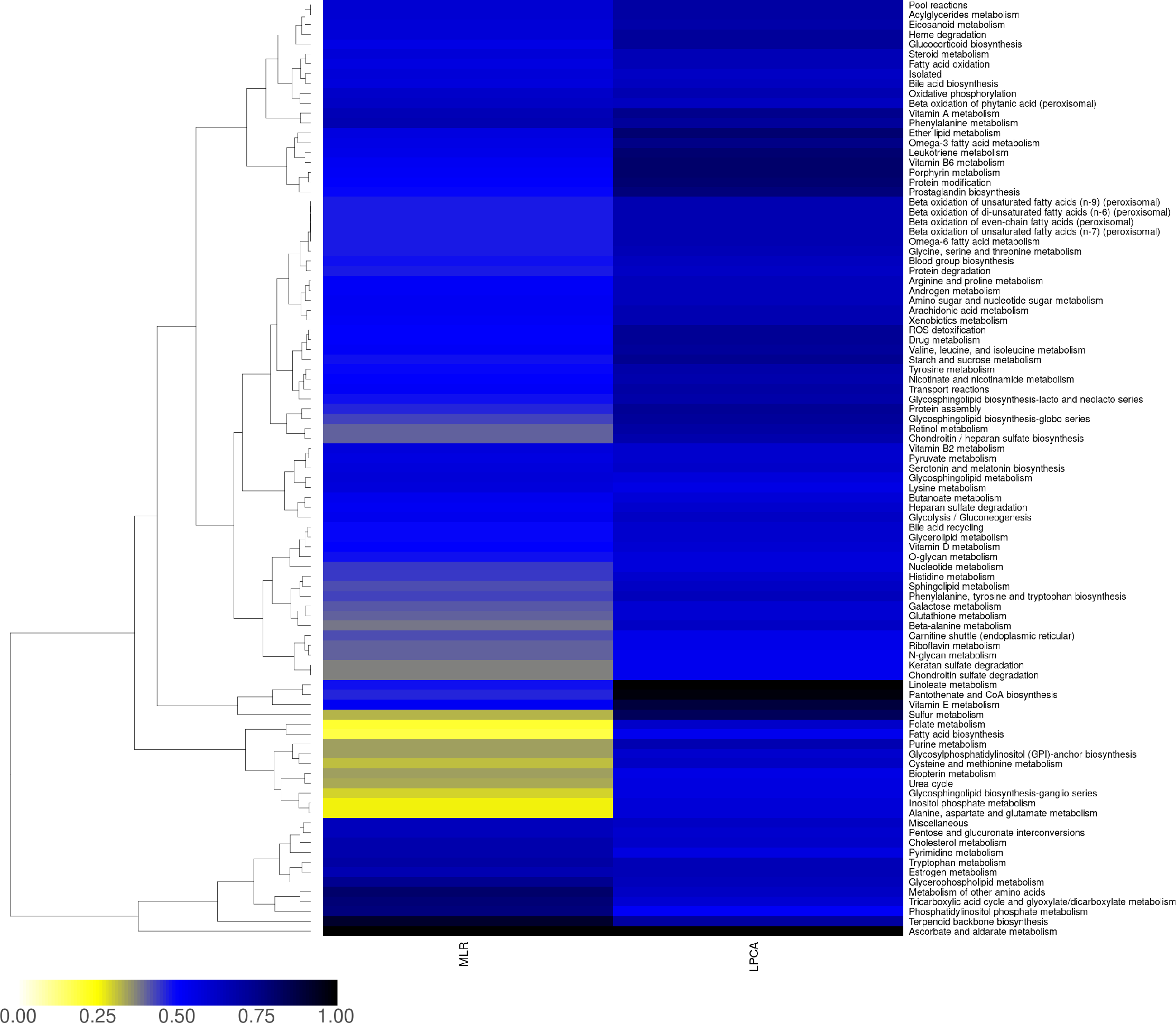
Impact of subsystems from LPCA and MLR for human reconstructions. MLR: contribution of subsystems to phylogenetic classification normalized to the maximum value. LPCA: subsystem-centric loadings normalized to maximum loading (refer to methods section for details)

### **Analysis of** 2943 Agora2-derived *Firmicutes* species

Finally, we applied LPCA to a subset (*Firmicutes*) of the Agora2 dataset, after creating a binary reaction matrix, containing 5267 differential reactions from 2943 species-specific GSMMs. The three main orders (*Bacillales*, *Eubacteriales*, and *Lactobacillales*) formed subclusters regardless of the applied method (Fig 7). Incomplete subsystem assignments within the GSMMs prevented the determination of meaningful subsystem-centric loadings. This underscores the significance of accurate subsystem assignments when utilizing LPCA for GSMMs.

**Fig 7.**
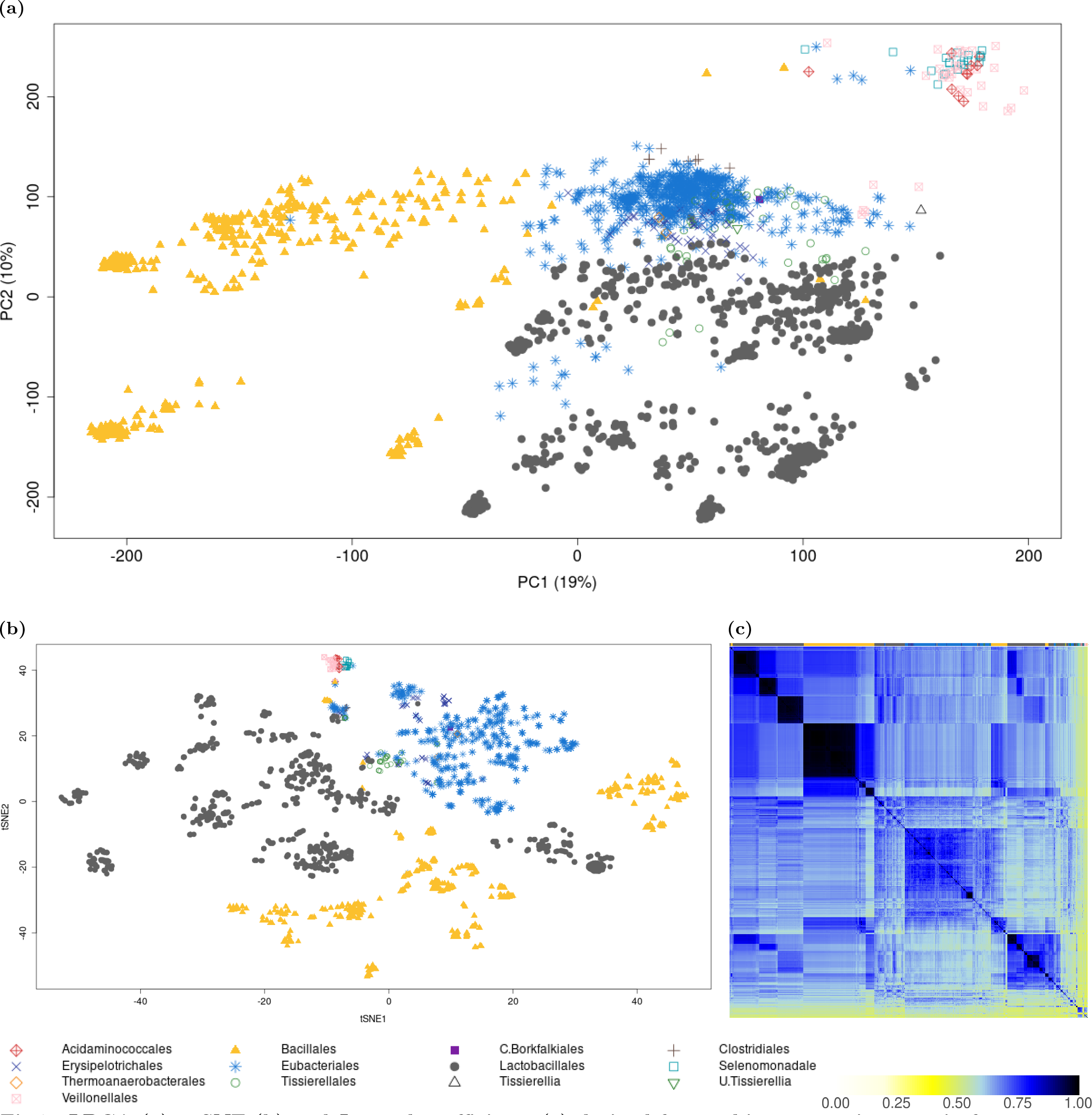
LPCA (a), t-SNE (b) and Jaccard coefficients (c) derived from a binary reaction matrix from 2943 *Firmicutes* species-GSMMs. In panels (a) and (b), points represent individual GSMMs, with different genera indicated by distinct symbols and colors. The top row in panel (c) uses these same colors to indicate the corresponding genera. The clustering of rows and columns in panel (c) was performed using the default hierarchical clustering settings (refer to methods section for details)

We assessed the run time for LPCA, t-SNE, and Jaccard coefficients. While t-SNE and Jaccard coefficients finished in 0.31 h and 1.26 h, respectively, LPCA took 20 h on an “AMD EPYC 7542 32-Core Processor”, 96 cores, 406 GB RAM. While LPCA offers valuable insights into GSMMs at the subsystem and reaction levels, its longer runtime may pose challenges for huge datasets and could benefit from further optimization.

## Conclusion

Here we introduced LPCA to simultaneously compare and analyze multiple GSMMs to identify similarities and differences in metabolic capabilities across multiple species and strains. LPCA extends standard PCA to binary data sets. Thus, it allows us to analyze the presence or absence of biochemical reactions across GSMMs. Our approach not only confirmed the established phylogenetic relationships but also demonstrated the robustness of LPCA in delineating clustering patterns. Utilizing LPCA on the set of differential reactions across GSMMs provides distinct advantages:

1. Enhanced Clade Discrimination: Unlike PCA relying on simulated growth rates from varied environmental conditions, LPCA exhibited clear separation of distinct clades. This method mitigates bias by solely assessing reaction presence in GSMMs, rather than subjectively selecting environmental conditions for growth rate simulations.
2. Subsystem Identification: In contrast to t-SNE and Jaccard coefficients, our LPCA approach provides precise information on drivers that govern separation and enables an efficient subsystem analysis by grouping reactions based on their loading values. This may also support curation efforts and metabolic subsystem analysis.
3. Comparison to MLR: While MLR can identify driving factors, LPCA offers a significant advantage by simultaneously identifying potential subclusters. These subclusters may indicate metabolic variations that could be overlooked when relying solely on the initial phylogenetic classification. Additionally, the direction of reaction- or subsystem-centric loadings can be determined with LPCA, which is not possible with MLR.
4. Transcriptome-based Equivalence: Our findings showcased the ability of LPCA to recover cluster patterns observed in transcriptome-based PCA plots solely from genome-scale metabolic reconstruction and/or GSMMs.

Here, our focus was on grouping reactions by standard biochemical subsystems such as glycolysis, etc.. However, LPCA can be extended to group reaction-centric loadings by any other pathways of interest. This expansion could complement recently developed tools designed to facilitate metabolic pathway analysis [48, 49]. In addition to other methods for feature selection from omics datasets [50–52], PCA loadings have been established as an effective method for this purpose, facilitating the identification of biologically significant features with high variance across different conditions or phenotypes [53]. LPCA loadings could be used in a similar way with respect to GSMMs. While LPCA is limited to binary datasets, our study demonstrates its effectiveness with binary reaction datasets derived from GSMMs, highlighting its potential to guide further research and pathway analysis in GSMMs. By identifying reactions with high LPCA loading values, we have pinpointed those that play pivotal roles in GSMMs, suggesting targets for further experimental and computational investigation. Integrating this approach with prior to other omics analyses could ultimately provide a more comprehensive understanding of metabolic pathways.

## Author’s contributions

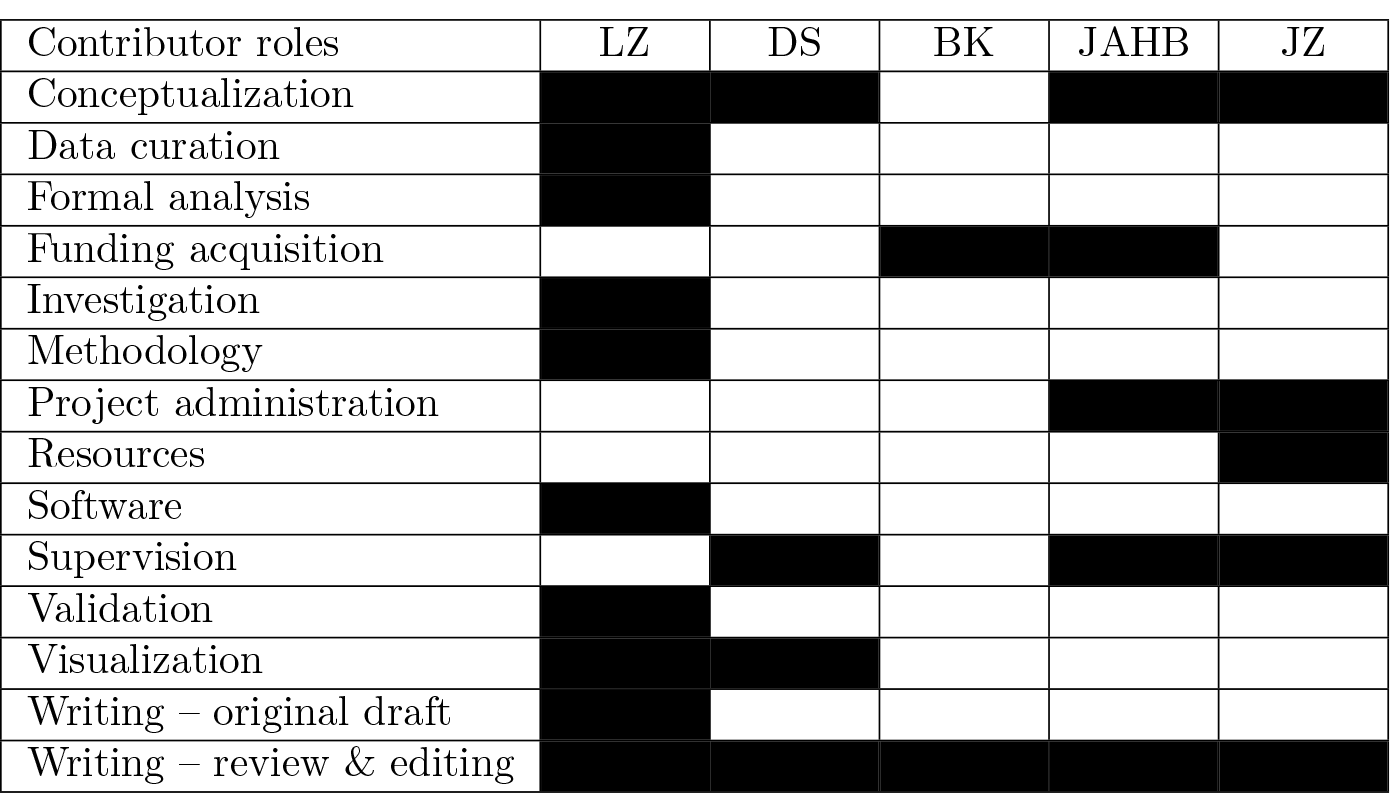

## Acknowledgment

LZ, BK, and JAHB are employees of Baxalta Innovation GmbH. Employees of Baxalta Innovations GmbH may be owners of stock and/or stock options. DS and JZ are employees of the University of Vienna. The presented work was partly funded by Baxalta Innovations GmbH. The funder had no role in the study design. The funder approved the decision to publish.

## Data availability

All data and codes underlying this article are freely available in a GitHub repository, at https://github.com/LeoZ93/lpca4gsmm.

## Supplementary Information

Fig S1 . LPCA (a), t-SNE (b) and Jaccard coefficients (c) derived from a binary reaction matrix from pan-reactions in 222 *Escherichia* GSMMs. In panels (a) and (b), points represent individual GSMMs, with different genera indicated by distinct symbols and colors. The top row in panel (c) uses these same colors to indicate the corresponding genera. Circles in panel (a) highlight clusters of *E. albertii* strains (blue), *E. fergusonii* strains (red), and a mixed cluster of *E. coli*, *S. dysenteriae*, *S. flexneri*, and Clades II to VIII (orange). Labeled arrows in panel (a) denote subsystem-centric loading vectors from LPCA (refer to the results and methods section for definitions). The clustering of rows and columns in panel (c) was performed using the default hierarchical clustering settings (refer to methods section for details)

Fig S2 . PCA based on simulated growth rates from 222 *Escherichia* species-specific GSMMs across 570 different environmental conditions [9]. While *E.fergusonii* GSMMs could be well separated based on simulated growth rates, the remaining clades seem to be less separatable.

Fig S3 . Phylogenetic tree of 222 *Escherichia* species based on whole genomes. Genomes were obtained from Enterobase [30]. Phylogenetic relations were obtained from OrthoFinder [32] based on coding genes. *E.coli*, *E.albertii*, and *E.fergusonii* formed distinct clades.

Fig S4 . Reaction-specific loadings, grouped by subsystem from differential reactions. The reaction “ALAALAD” (Murein Biosynthesis) was found to be a major driving factor for separation. “PRCOA1” (Cholesterol degradation) was found to be incorrectly assigned to “Histidine metabolism” in the original GSMMs.

Fig S5 . LPCA scores (a) and t-SNE (b) results using the differential reaction dataset from *Escherichia* -GSMMs, performed 10 times. While LPCA scores showed reproducible clustering, t-SNE resulted in a more diffuse clustering.

Fig S6 . Phylogenetic tree of yeast-species taken from [24]. Whole-genome based phylogenetic comparison of strains results in more distinct separation of clades, compared to LPCA scores, t-SNE or hierarchical clustering based on Jaccard similarity.

Fig S7 . PCA of transcriptomes from (a) whole transcriptomes, (b) “Human1” metabolic genes, and (c) selected genes from context-specific reconstructions. While clustering between healthy and cancer tissue could be conserved based on whole transcriptomes (a) and metabolic genes (b), it was not possible based on the genes from the context-specific reconstructions (c).

Table S1 . Top-ranked subsystems driving cluster separation in 222 *Escherichia* strains in LPCA. Subsystems were ranked either according to the magnitude of the average loadings per subsystem for the first two principal components, avg *U^j^*, or according to the average magnitude of the loadings per subsystem, avg *U^j^*, see equations (2) and (3), respectively.

*†* not within the top five in this ranking

*‡* not present in the original set of subsystems

Table S2 . Reaction-centric loadings of the subsystem “Histidine metabolism”

*†* In the original models, “PRCOA1” was associated with “Histidine metabolism”. We reassigned “PRCOA1” to the newly created metabolic subsystem “Cholesterol degradation”. See text and Tab S1 for details.

